# Regulation of evidence accumulation by pupil-linked arousal processes

**DOI:** 10.1101/309526

**Authors:** Waitsang Keung, Todd A. Hagen, Robert C. Wilson

**Author notes:** First two authors share equal contribution. Department of Psychology, 1503 E University Blvd, Tucson, AZ 85719.

## Abstract

Integrating evidence over time is crucial for effective decision making. For simple perceptual decisions, a large body of work suggests that humans and animals are capable of integrating evidence over time fairly well, but that their performance is far from optimal. This suboptimality is thought to arise from a number of different sources including: (1) noise in sensory and motor systems, (2) unequal weighting of evidence over time, (3) order effects from previous trials and (4) irrational side biases for one choice over another. In this work we investigated these di.erent sources of suboptimality and how they are related to pupil dilation, a putative correlate of norepinephrine tone. In particular, we measured pupil response in humans making a series of decisions based on rapidly-presented auditory information in an evidence accumulation task. We found that people exhibited all four types of suboptimality, and that some of these suboptimalities covaried with each other across participants. Pupillometry showed that only noise and the uneven weighting of evidence over time, the ‘integration kernel’, were related to the change in pupil response during the stimulus. Moreover, these two different suboptimalities were related to different aspects of the pupil signal, with the individual differences in pupil response associated with individual differences in integration kernel, while trial-by-trial fluctuations in pupil response were associated with trial-by-trial fluctuations in noise. These results suggest that di.erent sources of suboptimality in human perceptual decision making are related to distinct pupil-linked processes possibly related to tonic and phasic norepinephrine activity.

## 1 Introduction

The ability to integrate evidence over time is a crucial component of perceptual decision making. This is true whether we are integrating visual information from saccade to saccade as we scan a scene, or integrating auditory information from word to word as we listen to someone talk. In recent years much work has been devoted to understanding how humans and animals perform evidence integration over short time scales (on the order of one second) in simple perceptual tasks [1, 2, 3, 4]. In a classic paradigm from this literature, known as the Random Dot Motion Task, participants are presented with a movie of randomly moving dots that have a weak tendency to drift in a particular direction (e.g. left or right) and they must decide which way the dots are drifting [5]. The optimal strategy in this task is to count, i.e. integrate, the number of dots moving to the left and right over the time course of the stimulus and choose the side that had the most dots moving in that direction. Amazingly, this optimal strategy can account for many of the qualitative properties of human and animal behaviour and neural correlates of integrated evidence can be found in several areas of the brain [2, 6, 3, 7].

Despite the ability of the optimal model to qualitatively account for a number of experimental findings, the quantitative performance of even highly trained humans and animals is suboptimal [1, 8]. This suboptimality is thought to arise from at least four different sources: (1) neuronal noise, (2) unequal weighting of evidence over time, (3) order effects from previous trials and (4) side biases.

The first source of suboptimality is neuronal noise. While the exact cause of neuronal noise is subject to debate [9, 10, 11, 8], it is thought that variability in neural firing impacts perceptual decision making in one of two ways. First, noisy sensory information reduces the quality of the evidence going into the accumulator in the first place [12, 1, 13, 14]. Second, noisy action selection causes mistakes to be made even after the integration process is complete [15, 16, 17].

The second source of suboptimality comes from the unequal weighting of evidence over time, which we call here the ‘integration kernel’. In particular, while the optimal kernel in most perceptual decision making tasks is flat — i.e. all information is weighed equally over time — a number of studies have shown that humans and animals can have quite suboptimal kernels. For example, in the Random Dot Motion Task, monkeys exhibit a ‘primacy’ kernel, putting more weight on the early parts of the stimulus relative to the later parts of the stimulus [4]. Conversely, in a slightly different integration task, humans exhibit the opposite ‘recency’ kernel, weighing later information more than early information [18, 19]. Finally, in some experiments this second source of suboptimality appears to be absent, with a ‘flat’ integration kernel being found in both rats and highly trained humans [1].

The third source of suboptimality reflects the tendency to let previous decisions and outcomes interfere with the present choice. Thus, when making multiple perceptual decisions, the current decision is influenced by the choice we just made, for example by repeating an action when it is rewarded and choosing something else when it is not, a reinforcement learning effect [15, 16, 8, 20], or simply repeating a choice regardless of the outcome associated with it, a choice kernel effect[8, 20, 21]. Such sequential dependence can be advantageous when there are temporal correlations between trials, as is the case in many reinforcement learning tasks [15, 16], but is suboptimal in most perceptual decision making tasks when each trial is independent of the past [22, 23, 8, 20].

Finally, the fourth suboptimality is an overall side bias where both humans and animals develop a preference for one option (e.g. left) even though that leads to more errors overall [20].

Evidence from a number of studies suggests that pupil-linked arousal processes, putatively driven by the locus coerulues norepinephrine system [24, 25, 26, 27, 28], are well placed to modulate all four of these different sources of suboptimality. With regard to noise, increased pupil response has been associated with a number of different cognitive processes such as effort, arousal, mood, attention and memory, all of which might influence noise [26, 29]. In the specific case of perceptual decisions, previous work suggests a role for pupil-linked arousal systems to modulate the overall neuronal noise, i.e. the signal-to-noise ratio of sensory cues, in the evidence accumulation process [30, 31]. With regard to kernel and side bias, pupil response has been associated with a change the ‘gain’ of other neural systems, which in turn is thought to modulate the strength of internal and external cognitive biases on decision making [26, 28, 32, 33, 34, 35]. In addition, recent empirical and theoretical work has also suggested that norepinephrine, a putative driver of pupil dilation, modulates the urgency of decision making in a sequential sampling task such that the higher the norepinephrine level, the more urgently a decision is made [36, 37, 38]. Taken together these studies point to the possibility of pupil-linked norepinephrine systems to modulate the integration kernel and side bias by changing the strength of pre-existing biases during the integration process. Finally, with regard to sequential effects, pupil changes have been related to how humans integrate relevant information from previous trials to infer uncertainty and expectation [21, 39, 40], suggesting a role for pupil-linked arousal systems in modulating sequential effects.

In this work we investigated all four sources of suboptimal perceptual decision making and their relationship to between pupil-linked arousal processes in a single task. By quantifying all four sources of suboptimality in the same task we were able to assess the relationships between the suboptimalities and determine the extent to which pupil-linked arousal processes were related to each.

## 2 Results

To study the effects of pupil response on evidence accumulation, we designed an auditory discrimination task based on the Poisson Clicks Task [1]. In this task, participants listened to two trains of clicks in the left and right ears, and were instructed to indicate which side they thought had more clicks (Figure 1a). Clicks in our task were generated according to a Bernoulli process, such that there was always a click every 50ms that was either on the left, with probability *p*_*left*_, or otherwise on the right. This process meant that the total number of clicks was always fixed at 20 clicks and the clicks occurred at a fixed frequency of 20 Hz. This generative process for the clicks represented a slight departure from [1], in which clicks were generated by a Poisson process with a refractory period of 20ms. The main reason for using a Bernoulli process was to simplify the subsequent logistic regression analysis for quantifying the different sources of suboptimality, without imposing too much a priori assumptions. To indicate this difference we refer to our task as the Bernoulli Clicks Task.

### 2.1 Psychometric and chronometric functions

108 participants each performed between 666 and 938 trials (mean 760.7) of the Bernoulli Clicks Task. Basic behaviour was consistent with behaviours in similar pulsed-accumulation tasks [1, 4]. Choice exhibited a sigmoidal dependence on the net difference in evidence strength, i.e. the difference in number of clicks between the right and left, Δclick (Figure 1b). A simple logistic regression of the form:

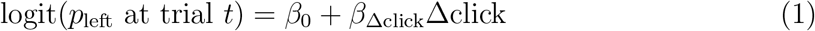

revealed a significant effect of Δclick (*β*_Δclick_ = 0.3500, *p* = 0.0001). Reaction times were also modulated by net evidence strength (Figure 1c) and linear regression of the form:

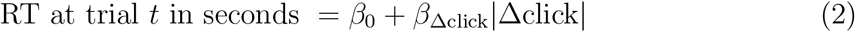

found a significant effect of the absolute value of Δclick on RT (*β*_Δclick_ = −0.017, *p* = 3.7 × 10^−13^). These results indicated that participants were faster and more accurate when the difference of number of clicks was large (easy trials), and less accurate and slower when that difference was small (hard trials).

**Figure 1:**
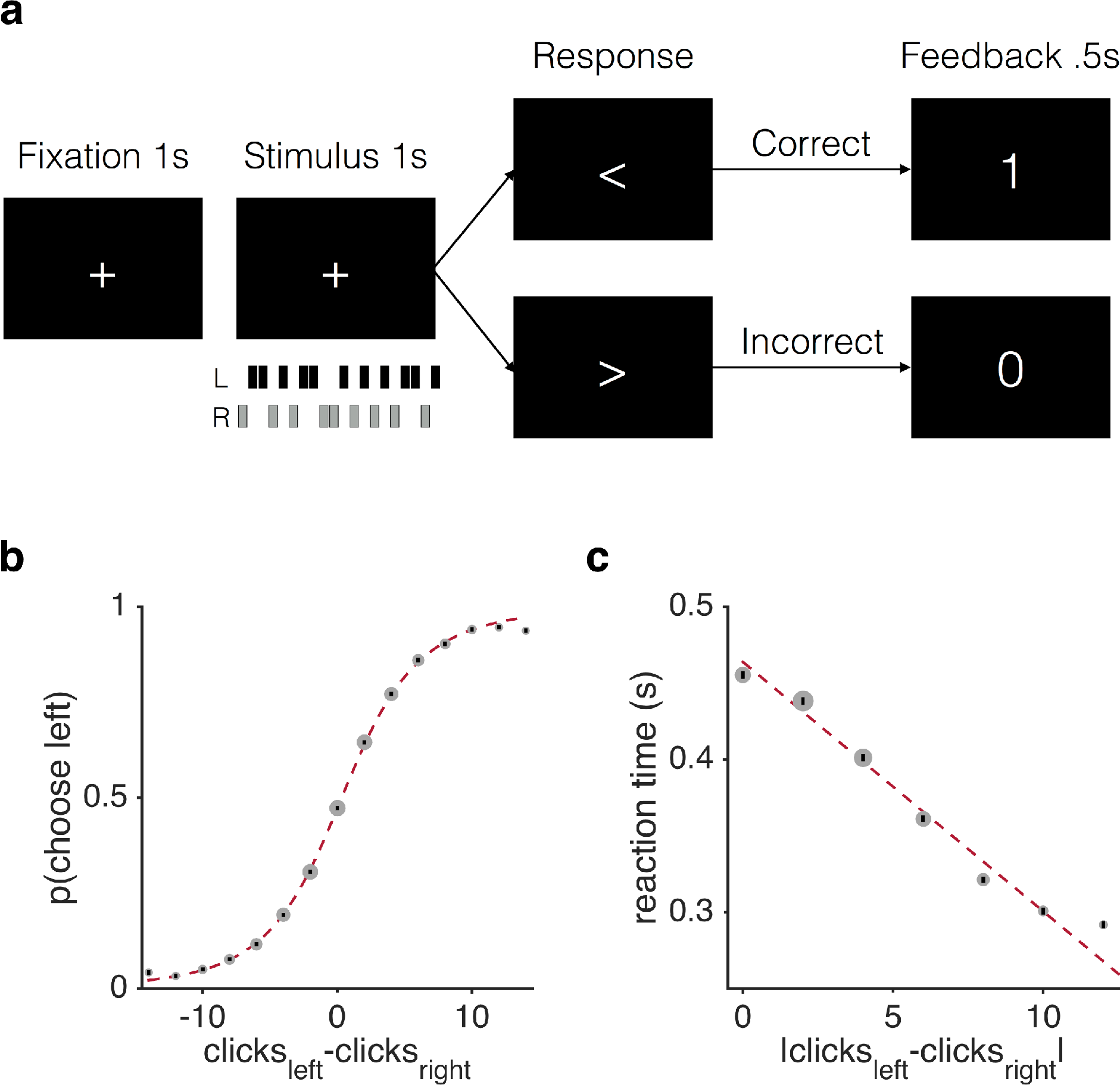
Basic behaviour of 108 participants. (a) Participants listened to a train of twenty clicks coming in either the left (L, black bars) or right (R, grey bars) ear for one second, and decided which side had more clicks. (b) Choice probability (probability of choosing left) showed sigmoidal relationship with difficulty (the difference in number of clicks between left and right). (c) Reaction times were higher on more difficult trials. Size of grey dots scaled by number of trials. All error bars (black bars) indicate s.e.m. across participants. Dotted red lines are fits with sigmoidal function and linear function respectively.

### 2.2 Humans exhibited all four suboptimalities in the Bernoulli Clicks Task

We used a logistic regression model to characterize the four different types of suboptimalities in human decision making in our task. This model quantified the impact of each click, the reinforcement learning and choice kernel effects from the five previous trials and the side bias on participants’ choices. In particular, we assumed that the probability of choosing left on trial *t* was given by

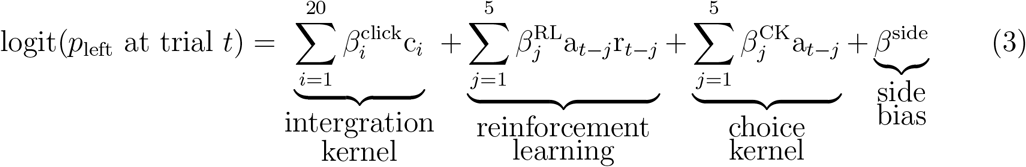

where c_*i*_ was the *i*th click (+1 for a left click and −1 for right), *a*_*t*−*j*_ was the choice made on the *t*-*j*th trial (+1 for a left choice and −1 for right), and r_*t*−*j*_ was the ‘reward’ on the *t*-*j*th trial (+1 for correct and −1 for incorrect). Therefore, a_*t*−*j*_r_*t*−*j*_ indicated the correct side on the *t*-*j*th trial (+1 when left was correct and −1 when right was correct). The relative effect of each of these terms on the decision was determined by the regression weights: 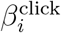 (the effect of each click), 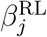 (the reinforcement learning (RL) effect, i.e. effect of previous correct side), 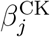 (the choice kernel (CK) effect, i.e. effect of previous choice) and *β*^side^ (an overall side bias).

**Figure 2:**
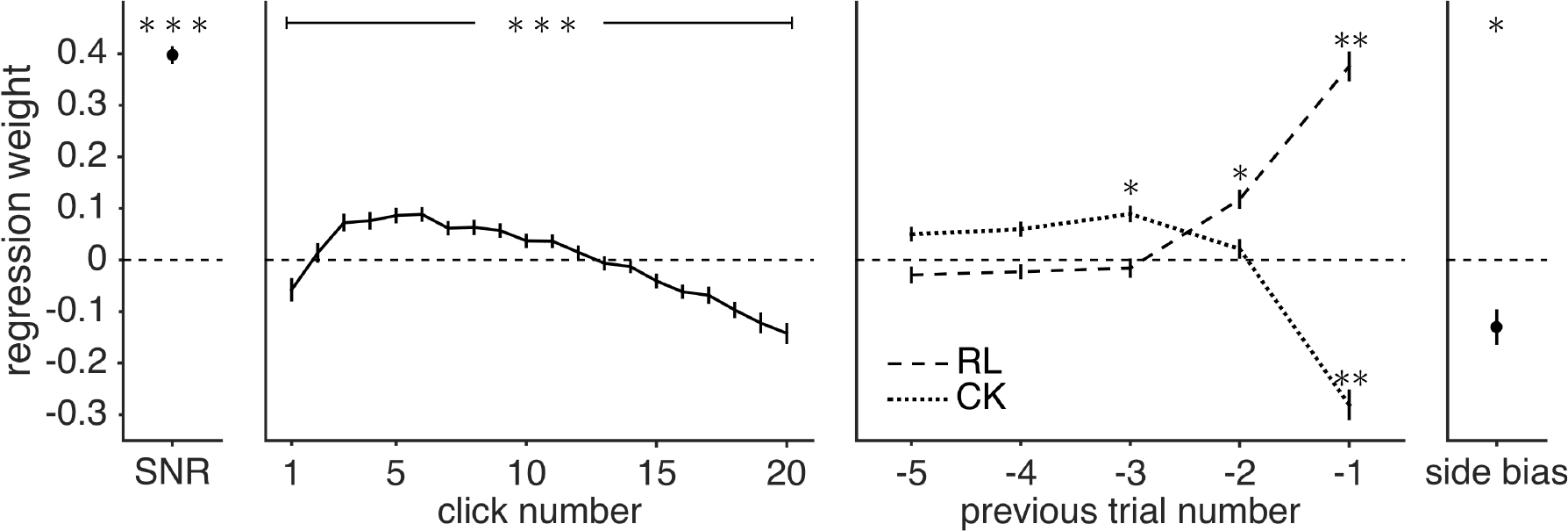
Regression model. (Leftmost) Mean of click weights is significantly above zero. T-test against zero, two-tailed, ***: FDR corrected for multiple comparisons *p* < 0.00001. (Second from left) Deviation of click weights from the mean has an uneven shape. repeated measures ANOVA, ***: *p* < 0.00001. (Second from right) Effect of previous trials: RL (the correct side in previous trial) positively predicts choice, indicating a reinforcement learning effect, while as CK (the choice made in previous trial) negatively predicts choice, indicating a alternating choice kernel. T-test against zero, two-tailed, **: FDR corrected for multiple comparisons *p* < 0.00001, Cohen’s d > 1; *: FDR corrected for multiple comparisons *p* < 0.00001, Cohen’s d > 0.5. (Rightmost) Side bias. T-test against zero, two-tailed, *: FDR corrected for multiple comparisons *p* = 0.0001. All error bars (black bars) indicate s.e.m. across participants.

Each of the four suboptimalities could be quantified using different parameters from this model (Figure 2). First, the signal-to-noise ratio (SNR), corresponding to suboptimality arising from neuronal noise, was quantified as the average weight given to all clicks 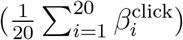. The higher the average click weight, the higher the SNR or equivalently, the lower the relative level of the noise. This average was significantly different from zero (*T*(107) = 27.65, two-tailed, FDR corrected for multiple comparisons *p* < 0.00001, Cohen’s d = 2.66) (Figure 2 leftmost panel), indicating that participants based their decision on (at least some of) the clicks and that each click increased the log odds of ultimately choosing that direction by about 0.4.

The second suboptimality, i.e. deviations from a flat integration kernel, was quantified as the deviation of the click weights from the average, i.e. 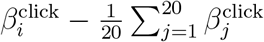 (Figure 2 second from left panel). Here we found that participants did not weigh all the clicks equally (repeated measures ANOVA, *F*(19, 2033) = 28.21, *p* < 0.00001, partial *η*^2^ = 0.21). This was not consistent with previous reports with a similar task where all clicks received equal weighting on average [1].

Sequential effects, the third suboptimality, were captured by the effects from previous trials. Specifically, the terms 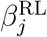 and 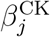 quantified the reinforcement learning (RL) effect (effect of past correct side) and choice kernel (CK) effect (effect of past choice) for the past five trials on the current choice. In line with earlier work [20], we found that previous trials had both significant RL and CK effects on participants’ choices (Figure 2 second from right panel). Notably, the positive RL regression weight demonstrated a positive reinforcement learning effect, in that participants tended to choose whichever side that was shown to be correct on the previous trial (*T*(107) = 14.40, two-tailed, FDR corrected for multiple comparisons *p* < 0.00001, Cohen’s d = 1.39). The negative CK regression weight indicated an alternating choice kernel — participants tended to choose the opposite of what they had chosen on the previous trial (*T*(107) = −10.45, two-tailed, FDR corrected for multiple comparisons *p* < 0.00001, Cohen’s d = −1.01).

Finally, the side bias was quantified by the intercept term *β*^side^ in the model (Figure 2 rightmost panel). This term quantified the extent to which a participant chose the left side on all trials regardless of which side was the correct side. Here we saw a significant right bias indicated by a significantly negative regression weight (*T*(107) = −4.12, two-tailed, FDR corrected for multiple comparisons *p* = 0.0001, Cohen’s d = −0.40).

### 2.3 Sequential effects and signal-to-noise ratio covary across participants

We then inspected how these suboptimalities correlated with each other across participants. We used a three way mixed ANOVA to inspect the effect of previous trials on both SNR and kernel shape. The three factors we investigated were: RL regression weights, CK regression weights, and time. The ANOVA was set up to investigate the effect of these three factors on the regression weights of clicks. In this ANOVA, the main effects of either RL or CK on kernel weights told us whether RL or CK correlated with the overall SNR. The interaction effect between either RL or CK and time on kernel weights told us whether RL or CK correlated with the kernel shape.

**Figure 3:**
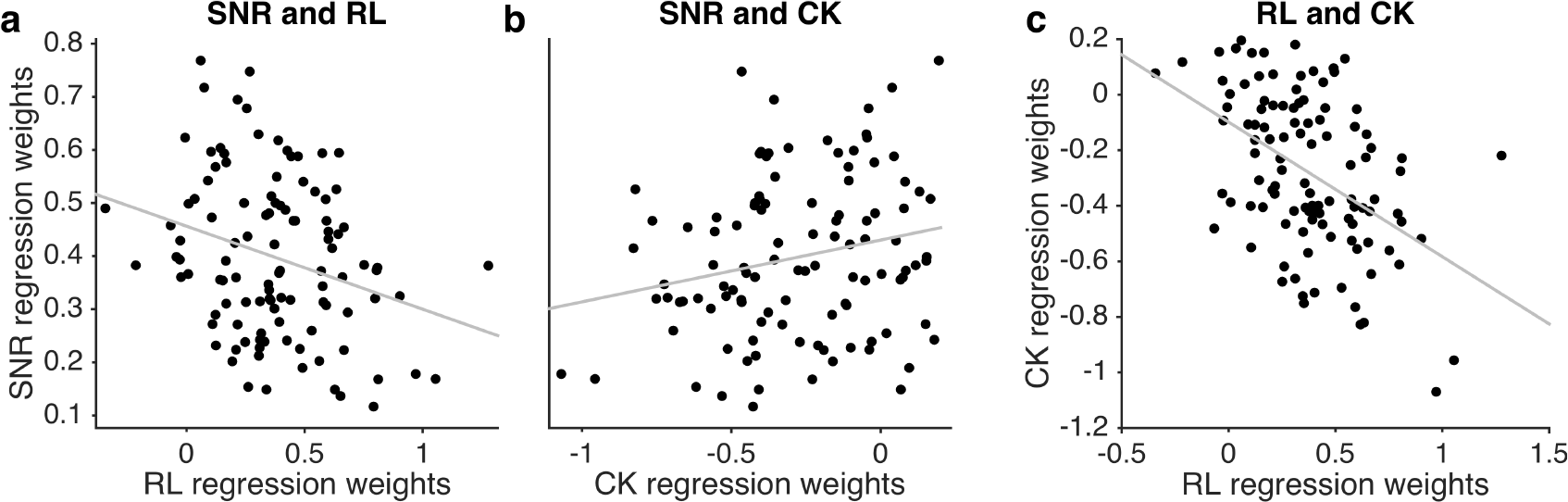
Interaction between suboptimalities across participants. (a) RL is significantly negatively correlated with SNR (*r* = −0.28, *p* = 0.003). (b) CK is significantly positively correlated with SNR (*r* = 0.22, *p* = 0.03). (c) RL is significantly negatively correlated with CK (*r* = −0.46, *p* = 5 × 10^−7^).

We found that both RL and CK had significant main effects on SNR (RL: *F*(1, 2080) = 50.66, *p* = 1.5 × 10^−12^, CK: *F*(1, 2080) = 56.82, *p* = 7.1 × 10^−14^), but not kernel shape (RL × time: *F*(19, 2080) = 0.80, *p* = 0.71, CK × time: *F*(19,2080) = 0.19, *p* = 0.99). Specifically RL is negatively correlated with SNR (*r* = −0.28, *p* = 0.003) (Figure 3a), while CK is positively correlated with SNR (*r* = 0.22, *p* = 0.03) (Figure 3b).

Importantly, since the RL effect was positive (Figure 2), i.e. participants tended to choose, on the current trial, whichever side was correct in the previous trial), a negative correlation indicated that participants who relied more on feedback from the previous trial tended to rely less on information on the current trial. Conversely, since CK effect was negative (i.e. participants tended to alternate their choices of sides between trials) (Figure 2), a positive correlation indicated that the more participants alternated their choices (i.e. relied on past choice history), again the less they relied on evidence from the current trial.

Together these results suggested a ‘subtractive’ effect between choice history and signal-to-noise ratio on the current trial - participants who rely more on history (RL and CK) tend to rely less on evidence from the current trial. This result could also be interpreted as that participants who were worse at making decisions based on evidence from the current trial tended to rely more on previous history. Interestingly, we also found a similar small but significant relationship between sequential effects and SNR at the within participant level (Supplementary Materials Section S.1).

In addition, we also saw a negative correlation between RL and CK across participants (*r* = −0.46, *p* = 5 × 10^−7^) (Figure 3c), which indicated that participants who relied more on past feedback also relied more on past choice (stronger alternating effect).

### 2.4 Individual differences in pupil change correlate with individual differences integration kernel

To examine the interaction between individual differences in pupil response and integration behaviour, we first computed the pupil diameter change during the presentation of clicks stimulus. We time-locked the pupillary response to the onset of the clicks stimulus, and averaged the pupil diameter within each participant. We then took the difference between the peak and the trough of the pupil diameter within the clicks stimulus, which we called the the ‘pupil change’ for each participant (Figure 4a). As shown by a median split in Figure 4b, there were considerable individual differences in the pupil change with some participants showing almost no change while others changed a lot during the stimulus.

To examine the relationship of pupil change with overall signal-to-noise ratio and integration kernel, we used a two way mixed ANOVA to compare the effects of pupil change(coded as a continuous variable) and time on participants’ regression weights from equation (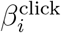s from equation (3)). If pupil change had an effect on the overall signal-to-noise ratio, we should see a main effect of pupil change on the regression weights. Conversely, if pupil change had an effect on the integration kernel, we should see a significant interaction effect between pupil and time on regression weights. Only the interaction effect was significant (interaction *F*(19, 2014) = 2.225, *p* = 0.0018, partial *η*^2^ = 0.02; main effect *F*(1, 106) = 2.761, *p* > 0.05). Moreover, these results were robust to a number of different assumptions in the analysis such as the size and location of the window size for computing pupil change (Supplementary Materials Section S.4), and whether we performed the analysis on the raw regression weights or on the top two principal components (Supplementary Materials Section S.3). Taken together these findings suggested that individual differences in pupil change affected the shape of the integration kernel but not the overall signal-to-noise ratio (illustrated using a median split in Figure 4c left two panels).

**Figure 4:**
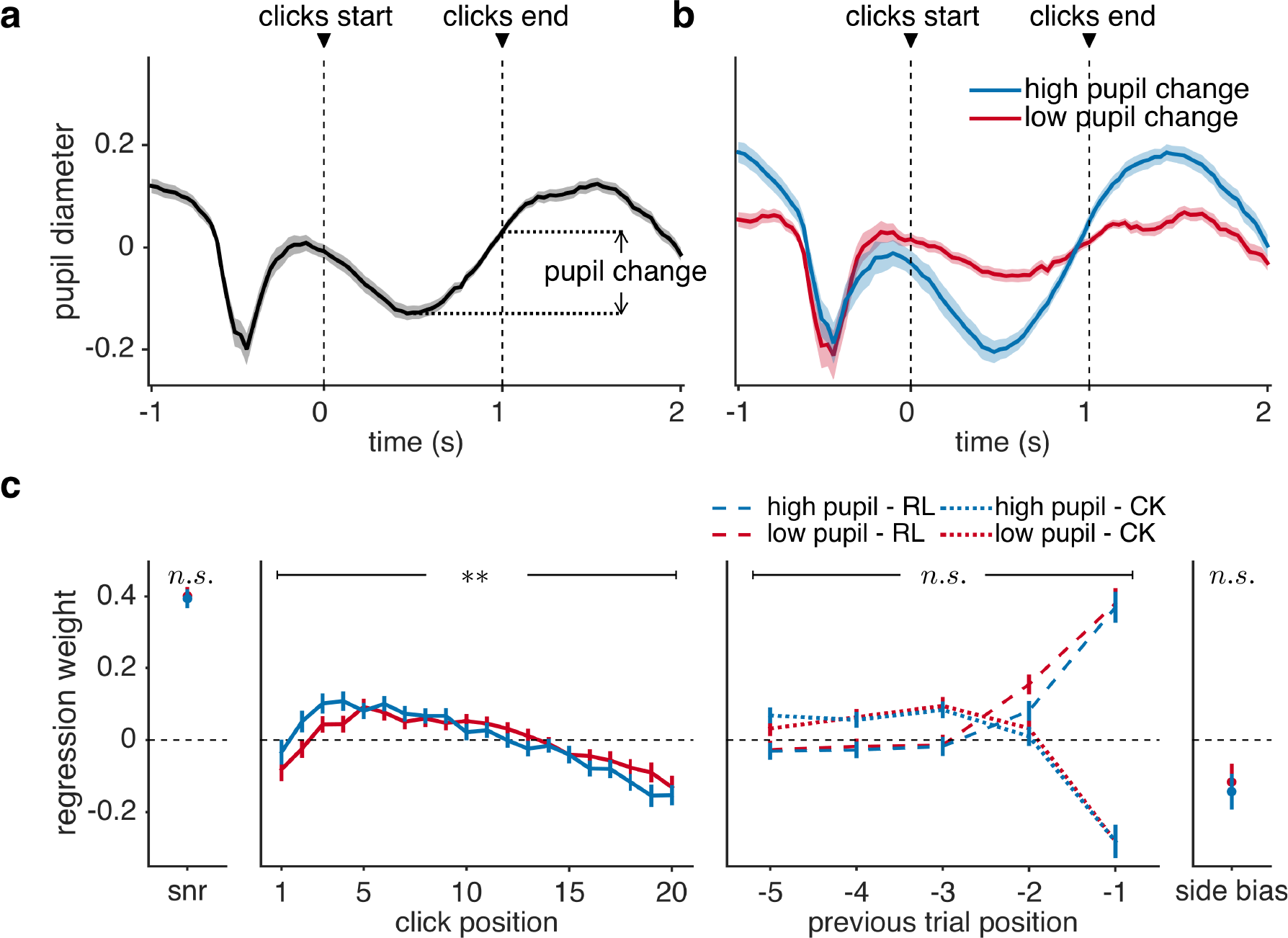
Interaction between pupil change and integration behaviour across participants.(a) Pupil diameter time-locked to the onset of clicks, averaged within participants across trials and then across participants. All shaded areas indicate s.e.m. across participants.(b) Averaged pupil response across participants split into two groups — high (blue) vs. low (red) change in pupil response. All shaded areas indicate s.e.m. across participants.(c) Regression weights averaged across participants split into high vs. low pupil change groups for visualization. (Leftmost) Mean regression weight showed no change across groups. (Second from left) Pupil had a significant interaction effect with time on regression weights. Two way mixed ANOVA, ** *p* = 0.0018. (Second from right) Effect of previous trials showed no differences across groups. (Rightmost) Side bias showed no change across groups. All error bars indicate s.e.m. across participants.

To understand which click weights were driving this interaction effect we performed a correlation analysis between individual differences in the regression weights for each click and individual differences in the pupil change. These post hoc tests suggested that the main change occurred in the second and third clicks, whose weights were increased in participants with high pupil change. Specifically, pupil change was significantly correlated with 2nd (*r* = 0.29, FDR corrected for multiple comparisons *p* = 0.02) and 3rd (*r* = 0.30, FDR corrected for multiple comparisons *p* = 0.02) kernel weights (Figure S2).

To examine the relationship between pupil change and sequential effects and side bias, we looked at the correlation between pupil change and regression weights for the previous trials (RL and CK) and side bias. We found no significant relationship between pupil change and either sequential effects (absolute correlation *r* < 0.16, FDR corrected for multiple comparisons *p* > 0.05) or side bias (correlation *r* = −0.01, FDR corrected for multiple comparisons *p* > 0.05) (illustrated using a median split in Figure 4c right two panels)

These results combined suggest that individual differences in pupil change were associated with individual differences in only one of the four suboptimalities, the kernel of integration such that participants with larger pupil change had more uneven integration kernels.

### 2.5 Trial-by-trial variability in pupil change correlates with trial-by-trial variability in signal-to-noise ratio

To quantify how trial-by-trial pupil change relates to the four suboptimalities in evidence accumulation, we modified the regression model (equation 3) to include interaction terms between clicks, previous trials, and trial-by-trial fluctuations in pupil:

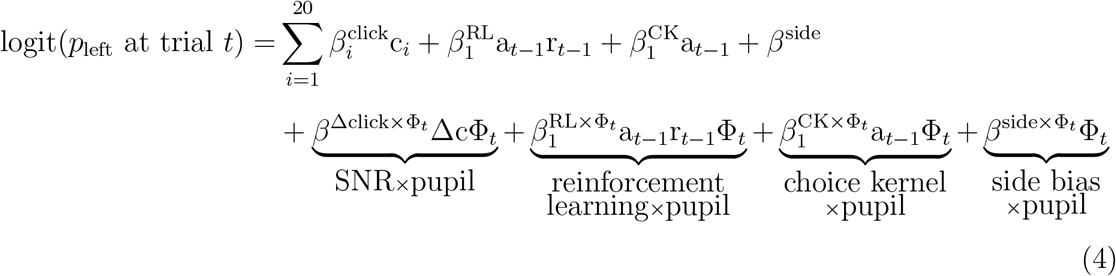

where Δc was number of clicks on the left minus number of clicks on the right, corresponding to the mean click regression weight in Figure 2, indicating the average signal-to-noise ratio. (It is also worth noting that here the interaction between side bias and pupil is equivalent to a main effect of pupil change - since the regressor indicating a side bias is all 1s.)

We found that trial-by-trial pupil change interacted significantly with Δclick after correction for multiple comparisons (*T*_107_ = −3.27, two-tailed, FDR corrected for multiple comparisons *p* = 0.0016, Cohen’s d = −0.31), but not with side bias, RL (previous correct) or CK (previous choice) (Figure 5).

**Figure 5:**
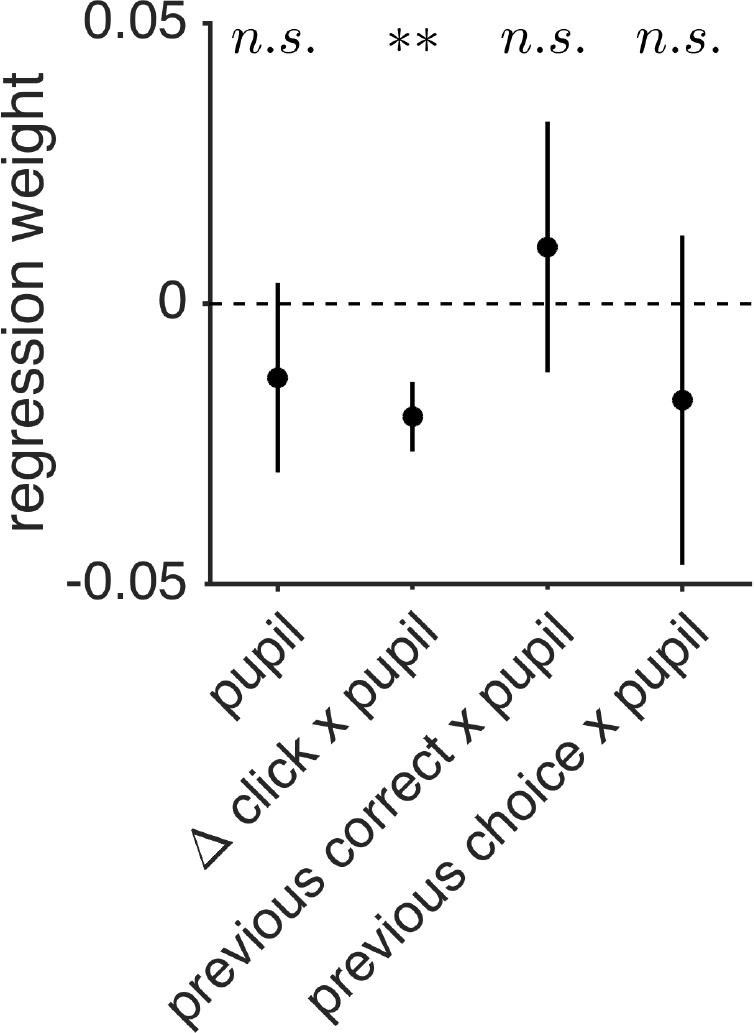
Trial-by-trial interaction between pupil change and integration behaviour. T-test against zero, two-tailed, **: FDR corrected for multiple comparisons *p* = 0.0016. All error bars (black bars) indicate s.e.m. across participants

We then tested whether there was an interaction between pupil change and integration kernel shape with a slightly modified version of equation (4):

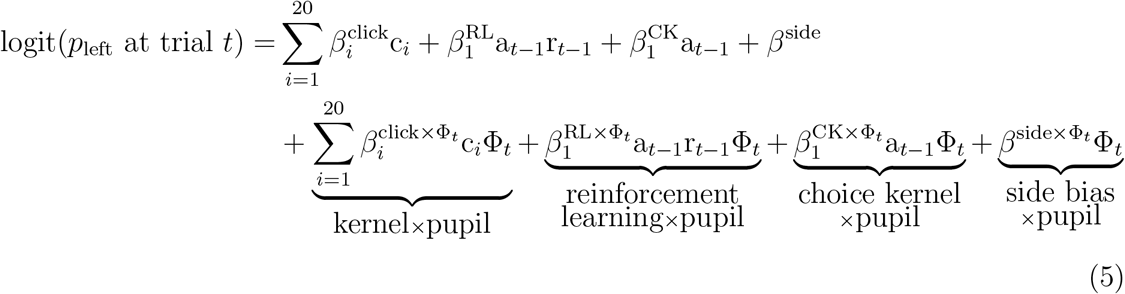

where Φ_*t*_ was the pupil change measure at trial *t*. The first four terms in this model were the same as equation (3), and the last four terms were the respective interaction terms of clicks (integration kernel), previous correct side (RL), previous choice (CK), and side bias with pupil change. With repeated measures ANOVA, we did not find a significant effect of time on 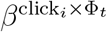 (*F*(19, 2033) = 0.72, *p* = 0.81), suggesting that pupil change did not modulate the integration kernel on a trial-by-trial level. We did not find a significant interaction effect between pupil and RL (*T*(107) = 1.78, two-tailed, FDR corrected *p* = 0.14), CK (*T*(107) = −0.78, two-tailed, FDR corrected *p* = 0.62), or side bias (*T*(107) = −1.63, two-tailed, FDR corrected *p* = 0.19) either. These results combined suggested that pupil change on a trial-by-trial level specifically modulated the overall signal-to-noise ratio, and not integration kernel or sequential effects.

## 3 Discussion

In this paper we investigated four sources of suboptimality in human evidence integration: neuronal noise (as reflected in the signal-to-noise ratio), uneven integration kernel, sequential effects, and side bias, and their relationship with pupil diameter at the across-participants and within-participants level. We showed that all four types of suboptimality were at play in our perceptual decision making task. These included variance that could not be explained by another source, i.e. ‘noise,’ a predominantly ‘bump’ integration kernel, sequential effects in the form of a positive reinforcement learning (RL) effect (choosing the previous correct answer) and an alternation choice kernel (CK) effect (choosing the opposite of previous choice), and an overall side bias (Figure 2). In addition, across the population, participants with stronger sequential effects (RL and CK) tended to rely less on evidence from the current trial (smaller signal-to-noise ratio), and participants with one kind of sequential effects (RL) also tended to have the other kind (CK) (Figure 3). At the physiological level, two of the four suboptimalities were associated with pupil dilation, at the trial-by-trial and individual difference levels respectively. At the individual difference level, only the integration kernel was associated with pupil change, with a more uneven profile of integration being associated with larger pupil change during stimulus presentation (Figure 4). Conversely, at the trial-by-trial level only noise was associated with pupil change, with a smaller signal-to-noise ratio being associated with larger pupil change on that trial (Figure 5). Our work adds to a growing literature on the suboptimalities in evidence accumulation and perceptual decision making and their relationship with pupil dilation. In the following we discuss the implications of our behavioral and pupillometric findings.

### 3.1 Behavioral findings

At the behavioral level, our findings are consistent with a number of previous results showing the presence of noise in the integration process [1, 8], uneven weighting of in-03 formation over time [4, 18, 19], and the presence of order effects [23, 21, 20] and side biases [1]. In addition, by running our task in a large number of participants (something not traditionally done in the animal literature), we were able to expose the relationships *between* the suboptimalities at the individual difference level. Intriguingly this analysis suggests an antagonistic relationship between the use of information from the past trial (i.e. RL and CK effects) and processing of the current stimulus, as reflected in SNR. Such an effect may reflect a kind of compensatory process in low performers. That is, people who are less able to process the stimulus correctly (low SNR) may rely more on sequential effects to (either explicitly or implicitly) try to compensate. While such a strategy is not adaptive for this task, this approach would pay off if there was autocorrelation in the task.

In addition to the correlations between suboptimalities, one unexpected behavioral finding was the *shape* of the uneven integration kernel. Specifically the ‘bump’ kernel where clicks in the middle are weighed more than those at the beginning or the end. This contrasts with previous work on perceptual decision making from Brunton and colleagues [1], who found that the integration kernel of rats and well-trained humans was flat, Yates and colleagues [4], who showed a purely primacy driven integration kernel in monkeys, and several studies that showed humans have a recency kernel [18, 19]. Given this difference in results, one obvious question is whether the bump kernel is a genuine feature of the integration process or some artifact of either the analysis pipeline or the task?

With respect to the analysis, one possibility is that the bump may result from a mixture of subjects with primacy and recency kernels which average together to form the bump. To test this we categorized the integration kernel for each participant intooneofthe following four shapes: bump, primacy, recency, and flat (for categorization method, see Supplementary Materials Section S.8). All 108 of these are plotted in Supplementary Figure S14. From here it is easy to see that a large number of subjects (49%) exhibit the bump kernel. This suggests that at least on the level of individual participants, the bump kernel is a feature of the integration process, and not just a artifact of averaging. Of course, the possibility remains that this bump kernel is a result of mixing a primacy and recency kernels *within subject* (e.g. some trials have primacy kernels, and some have recency). More detailed modeling work will be needed to tease these interpretations apart.

With respect to the task, another possible cause for the bump kernel comes from the number of clicks in each stimulus being fixed. This fixed number of clicks in each stimulus means that an ideal observer, who is aware that there are only 20 clicks in each stimulus, could safely stop integrating clicks if the excess number of clicks favoring on one side exceeds the number of remaining clicks. That is, by fixing the number of clicks, we may be implicitly favoring a bounded integration process (with a collapsing bound). Such bound crossing would cause the later clicks to be down-weighted on average as we see in the later part of the bump profile. Bound crossing would not, of course, account for the initial rise in weights for the bump profile, which would need some additional mechanism (perhaps a recency effect combined with a bound) to explain. Incidentally, this account would fit with the recency bias found in perceptual categorization in previous studies [18, 19]. An important direction for future research would be to test whether this account fully explains the bump profile, both with more detailed modeling in addition to more experiments in which the total number of clicks in each stimulus is not fixed.

### 3.2 Physiological findings

At the physiological level, our results add to a rapidly growing literature on the relationship between pupil dilation and decision making. In particular, this literature has reported associations between pupil dilation and a number of suboptimalities including: noise [21, 18], reinforcement learning effects [39], choice kernel effects [21] and pre-existing biases [28, 33]. Ours is the first to examine the relationship between pupil dilation and all of these suboptimalities in a single task, as well as being the first to look at the relationship between pupil dilation and the shape of integration kernel. Below we situate our results with respect to this previous literature considering each of the suboptimalities in turn.

### 3.2.1 SNR and pupil

With regard to SNR, we found that increased pupil change is associated with lower SNR on each trial. This finding is consistent with much of the previous literature. For example, in the Dot Motion paradigm, Murphy and colleagues showed that trial-by-trial variability in the evidence accumulation process was associated with increased pupil dilation [31]. Likewise in other perceptual decision making tasks several authors have observed an association with increased pupil dilation and noise in behavior [21, 18]. Outside of perceptual decision making, Jepma and colleagues observed the same relationship between pupil and decision noise in a reinforcement-learning based explore-exploit task [41].

Of course, while the finding that trial-to-trial pupil dilation is associated with trial-to-trial behavioral variability is robust across studies, exactly what this finding means is open to interpretation. In this paper we have related it to signal-to-noise ratio, with the interpretation that changes in pupil reflect change in SNR which causes poor performance. If one takes pupil as an index of activity in the locus coeruleus, our interpretation is consistent with the Adaptive Gain Theory of norepinephrine function, such that increased LC activity causes more variability in behavior via changes in neural gain [?, 42].

An alternate interpretation, put forth by Urai and colleagues [21], is that pupil reflects subjective uncertainty and that participants are more uncertain on trials in which they perform poorly. In this interpretation the direction of causality is reversed: it is poor performance that leads to changes in pupil, via its effect on uncertainty (which, incidentally, may also be related to LC [43]). Distinguishing between these accounts, which predict almost identical relationships between pupil and behavioral variability, will be difficult with correlational experiments such as ours, and future work using pharmacological and other causal interventions will be necessary to determine the direction of the relationship between pupil (putatitvely LC) and noise.

### 3.2.2 Sequential effects and pupil

In contrast to our result showing no relationship between pupil and the reinforcement learning and choice kernel, a number of studies have found relationships between pupil and sequential effects. For example, Nassar and colleagues showed that both baseline pupil and pupil change modulates how information from previous trials affect current choice [39], a result which was recently replicated in a different version of the task [40]. Similarly, in a perceptual decision making task, Urai and colleagues [21] showed that pupil dilation on the previous trial modulated the extent to which that trial influenced the current choice.

One possible cause of the difference between our results and this previous work is the overall magnitude of the sequential effects in the respective tasks. Specifically, in our task the sequential effects were small, with the combined effect of reinforcement learning and choice kernel equating to about 2 clicks, or 10% of the variance in the response. Conversely, in [21] the previous trial effects account for almost 100% of the variance when evidence on the current trial is weak. Likewise in [39] and [40] successful performance of the tasks *required* the use of sequential effects and so the sequential effects observed were huge. This difference in overall magnitude of the sequential effects could simply have made modulation of these sequential effects by pupil too small to observe in our task.

Another possible cause of the difference in results is the timing of the pupil signal that we focused on. Specifically, our task was optimized to look at pupil during presentation of the click stimuli and not at pupil at other points in the task, such as baseline pupil or pupil following the choice and feedback, which can have very differnet behavioral and computational correlates [44]. This difference in timing is especially important for the Urai et al. results [21] where the pupil signal modulating sequential effects was computed 250 ms before feedback, which is at least 2800 ms after stimulus onset. Such a time lag would be well into the inter-trial interval and possibly even the next trial in our task, making the corresponding pupil signal hard for us to compute. Indeed, when we looked at the signal at these later times, there was no association between pupil and sequential effects (Supplementary Materials Section S.5). Clearly, future experiments with additional delays will be necessary to determine whether pupil does or does not modulate sequential effects in our task.

### 3.2.3 Other biases and pupil

A number of other authors have related individual differences in pupil dilation to a number of other biases including risk aversion [45], learning styles [28], and the framing effect [33]. While these biases are not directly related to integrating evidence over time, the more general point that individuals with large pupil change have more bias across a range of tasks is consistent with our result that individual differences in pupil change modulate the integration kernel. In particular, we find that people with greater pupil change show more deviation (that is more bias) from the ideal, flat, integration kernel. Taken together these results suggests that, at least some, deviations from optimality are modulated by pupil, possibly via its association with LC.

### 3.2.4 Difference between between- and within-participant results

More generally, the difference between the between- and within-participant pupil results is intriguing. On the one hand, individual differences in pupil change correlate with kernel shape, while on the other trial-by-trial fluctuations in pupil change correlate with SNR. Why exactly would the individual differences and trial-by-trial correlates of the *same* signal be so different?

One possibility, originally raised in [28], is that these slightly different measures of pupil diameter — individual differences vs trial-by-trial fluctuation — may represent different neural measures, with the average of pupil dilations representing baseline or tonic locus coeruleus (LC) signals, and the trial-to-trial fluctuations of pupil reflecting transient, or phasic, LC firing. At the individual difference level, Eldar and colleagues [28, 33] have suggested that mean pupil response within a participant is a measure of tonic LC activity. In contrast, at the trial-by-trial level, work in monkeys and in mice has suggested that moment-to-moment pupil diameter changes track phasic LC firing [24, 25]. Applying these interpretations to the present findings suggests that tonic LC activity changes the kernel of integration while phasic LC decreases the signal-to-noise ratio.

The interpretation that tonic LC modulates the integration kernel between participants is consistent with previous work showing that individual differences in pupil change correlate with individual differences in susceptibility to a variety of cognitive and decision biases [28,33]. Importantly, theoretical work has shown with biophysically based neural network model that high tonic LC activity acts to amplify attractor dynamics, essentially causing the storage of impulsive decisions [38]. This can serve as a partial explanation for why our results revealed a positive correlation between individual pupil (proxy for tonic LC activity) and early kernel weights. Furthermore, empirical work has shown that pupil-linked arousal (associated with LC-NE activity and neural gain) is related to time-varying changes in the decision bound [36]. Specifically, higher pupil was found to reflect a stronger urgency signal (lower decision bound). While this relationship between pupil and urgency was only found for the case in which participants faced a decision deadline, an effect on decision bound could potentially offer an explanation for the relationship between pupil and the shape of the integration kernel. Clearly, more detailed modeling and experimental work will be needed to test this hypothesis.

The interpretation that phasic LC, as indexed by trial-by-trial pupil change, modulates signal-to-noise ratio is consistent with a number of pupil findings as outlined above [31, 18, 21]. However, it is at odds with a number of findings from direct LC recordings in monkeys, where enhanced phasic LC is associated with *better* task performance [46]. Understanding these results in more detail, with experiments in animals and neuroimaging in humans, will be important if we are to fully understand that LC plays in these decisions.

## 4 Methods

### 4.1 Participants

188 healthy participants (University of Arizona undergraduate students) took part in the experiment for course credit. We excluded 55 participants due to poor performance (accuracy lower than 60%), and then another 25 participants due to poor eye tracking data (see Eye tracking section below). All participants provided informed written consent prior to participating in the study. All procedures conformed to the human subject ethical regulations. All study procedures and consent were approved by the University of Arizona Institutional Review Board.

### 4.2 The Bernoulli Clicks Task

Participants made a series of auditory perceptual decisions. On each trial they listened to a series of 20 auditory ‘clicks’ presented over the course of 1 second. Clicks could be either ‘Left’ or ‘Right’ clicks, presented in the left or right ear. Participants decided which ear received the most clicks. In contrast to the Poisson Clicks Task [1], in which the click timing was random, clicks in our task were presented every 50 ms with a fixed probability (*p* = 0.55) of occurring in the ‘correct’ ear. The correct side was determined with a fixed 50% probability. Feedback appeared 500 ms after response, followed by a 1 s fixation delay before the next trial.

Participants performed the task on a desktop computer, while wearing headphones, and were positioned in chin rests to facilitate eye-tracking and pupillometry. They were instructed to fixate on a symbol displayed in the center of the screen, where response and outcome feedback was also displayed during trials, and made responses using a standard keyboard. Participants played until they made 500 correct responses or 50 minutes of total experiment time was reached.

### 4.3 Behavioural analyses

We modeled the choice with logistic regression using equation (3). In particular we assumed that the probability of choosing left on trial *t*, is a sigmoidal function of the impact from each click, the impact from five previous trial correct sides, the impact from five previous trial choices, and an overall side bias. In this model, by giving the *i*th click its own weight, we could account for the overall integration kernel.

### 4.4 Eye tracking

A desk-mounted EyeTribe eye-tracker was used to measure participants’ pupil diameter from both eyes at a rate of 30 samples per second while they were performing the behavioural task with their head fixed on a chin rest. Pupil diameter data were pre-processed to detect and remove blinks and other artifacts. Pupil diameter was z-scored across entire experiment before analysis. For each trial, pupil response was extracted time-locked to the start of the trial (Figure 4a). Change in pupil response was computed as the difference between the peak diameter and the minimum diameter during the 1s following trial onset. Pupil response measurements in which more than one-third of the samples contained artifacts were considered invalid and excluded from the analysis. Only participants with at least 200 valid trials were included in analysis (*n* = 108).

### 4.5 Across participants pupil analysis

For each participant, we took the mean pupil response across trials and computed the change in pupil diameter as described in previous section. We then compared this pupil change measurement with regression weights from equation (3). Specifically, we performed a two way mixed ANOVA in which pupil change is a between subject variable, time is a within subject variable, and regression weight is the dependent variable. We inspected the main effect of pupil change, which informed whether the average regression weight changed with pupil change across participants. We also inspected the interaction effect between pupil change and time, which informed whether pupil change modulates the effect of time on regression weights (i.e. the integration kernel).

### 4.6 Trial-by-trial within participants pupil analysis

For each trial, we took the pupil response and computed the change in pupil diameter. We then modeled participants’ choices with the logistic model in equations (4) and (5) to parse out trial-by-trial effects of pupil on integration. The first three terms in both equations were similar to equation (3). But in addition, we assumed that choice was also a function of the interaction between trial-by-trial pupil change and clicks, previous correct side, and previous choice.

### 4.7 Statistics

All data analyses and statistics were done in MATLAB and R. Repeated measures ANOVA, two way mixed ANOVA, and the corresponding post hoc tests done in R. All other analyses and statistical tests done in MATLAB.

## Author contributions

W. Keung did the data analyses. T. Hagen collected and preprocessed the data. T. Hagen and R. Wilson designed the experiment. W. Keung and R. Wilson wrote the manuscript. All three authors contributed to interpretation of the results and critical discussion.

## Acknowledgments

We thank Maxwell Alberhasky, Chrysta Andrade, Daniel Carrera, Kathryn Chung, Michael de Leon, Zamigul Dzhalilova, Asha Esprit, Abigail Foley, Emily Giron, Brittney Gonzalez, Anthony Haddad, Leah Hall, Maura Higgs, Marcus Jacobs, Min-Hee Kang, Kathryn 533 Kellohen, Neha Kwatra, Hannah Kyllo, Alex Lawwill, Stephanie Low, Colin Lynch, Alondra Ornelas, Genevieve Patterson, Filipa Santos, Shlishaa Savita, Catie Sikora, Vera Thornton, Guillermo Vargas, Christina West, and Callie Wong for help in running the experiments.

## Data availability

The data sets generated and analysed during the current study are available from the corresponding author upon reasonable request.

## Code availability

Experiment code was created with Psychtoolbox-3 and custom MATLAB code. All analyses were created with custom MATLAB and R code. All code will be uploaded to https://github.com/janekeung129/clicks-pupil.

## Supplementary Material

### S.1 Within participant analysis of interaction between sequential effects and signal-to-noise ratio (corresponding to Main Text Section 2.3)

Another way to investigate whether choice history enhances or diminishes the effect of current evidence is to inspect this interaction at a trial-by-trial level within participants. We adapted a technique for measuring consistency used by Cheadle and colleagues [18] – we measured the consistency *θ* between previous history (previous correct side and previous choice, representing reinforcement learning effect (RL) and choice kernel effect (CK) respectively) and the net difference in clicks on the current trial as the product between the two:

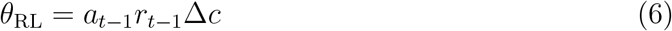

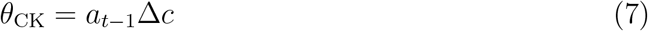

where Δ*c* was the net difference in clicks (each click coded as +1 for left or −1 for right),*a*_*t*−1_ was the previous choice (+1 for left or −1 for right), and *r*_*t*−1_ was the previous reward (+1 for correct and −1 for wrong), and *a*_*t*−1_*r*_*t*−1_ indicated which side (left or right) was correct in the previous trial (hereby referred to as previous correct side). If the previous correct side or previous choice and Δ*c* agreed in side (left or right), the consistency would be positive, and if not, the consistency would be negative. Because previous correct side and previous choice were always either +1 or −1, the magnitude of the consistency depended solely on Δ*c*, i.e. if the evidence on the current trial was strong and agreed with the previous trial, the consistency was highly positive, whereas if the evidence on the current trial was strong and disagreed with the previous trial, the consistency term would be highly negative.

We used this consistency measure as a “weight” by multiplying the term with the net difference in clicks, so that the weight of the current evidence is modulated by the consistency term. For example, in the case of *θ*_CK_, the regressor is computed as the following:

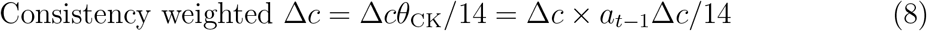

This regressor means that the impact of Δ*c* is enhanced (quadratically) when it agrees with previous choice, but is enhanced in the opposite direction if it does not agree with previous choice. We also normalized Δ*c* (by dividing it by the maximum absolute difference in clicks, which is 14) so that the Beta we estimate from this regressor will have the same unit with that of Δ*c*, for the purpose of comparing effect size. We added this interaction term in to the regression model in addition to the pure net difference in clicks term.

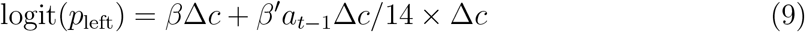

**Figure S1:**
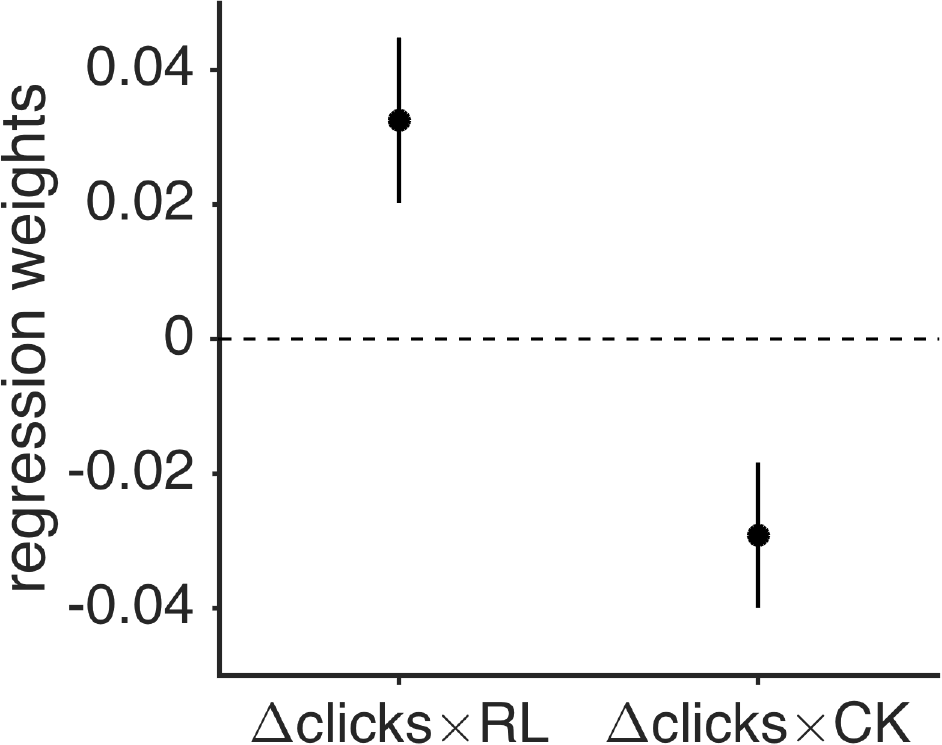
Trial-by-trial interaction between SNR and previous trial effects.

This consistency modulated Δ*c* came out significantly negative for CK (*b* = −0.029, FDR corrected for multiple comparsions *p* = 0.0075) and positive for RL (*b* = 0.032, FDR corrected *p* = 0.0087) (Figure S1). Since the previous history biases are positive for RL (reinforcement learning bias) and negative for CK (alternation bias), these results suggest that if Δ*c* is consistent with the reinforcement learning bias (previous correct side) it’s more highly weighted, and if Δ*c* is consistent with the alternating bias (i.e. not consistent with previous choice) it is again more highly weighted. This agrees with the individual differences analysis above, and suggests a multiplicative effect of previous trial bias on current trial on a trial–by–trial level. We do note that the effect sizes for both consistency-weighted Δ*c* are quite small, effectively amounting to about one-tenth of the effect of an average click.

### S.2 Scatter plots of correlation between early kernel weights and pupil (corresponding to Main Text Section 2.4)

We plot the scatter plots between pupil change and early (second and third) kernel weights below (Figure S2). Pupil change was significantly correlated with second (*r* = 0.29, FDR corrected for multiple comparisons *p* = 0.02) and third (*r* = 0.30, FDR corrected for multiple comparisons *p* = 0.02) kernel weights.

In addition, we removed a participant who may be an outlier (highlighted in red) in both correlations. The correlation results still held after removing the outlier: across participants, pupil change was significantly correlated with second (*r* = 0.29, FDR corrected for multiple comparisons *p* = 0.03) and third (*r* = 0.33, FDR corrected for multiple comparisons *p* = 0.01) kernel weights.

**Figure S2:**
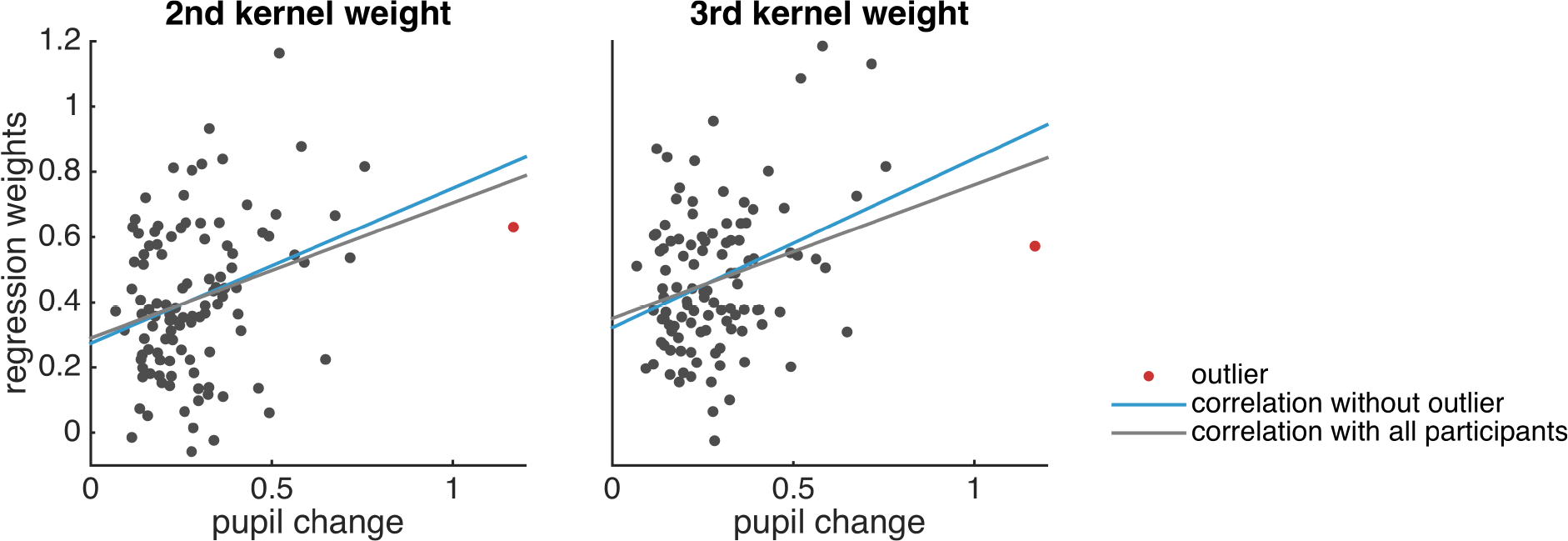
Scatter plots of second and third kernel weights and pupil change across participants. Grey line is least-squares line for all participants. Red dot is a potential outlier participant. Blue line is least-squares line for all particpants except the outlier.

### S.3 PCA on kernel weights (corresponding to Main Text Section 2.4)

We performed PCA on the kernel weights for dimensionality reduction. We showed the plot of the cumulative sum of the fraction of the total variance retained as the number of components increases (Figure S3a). We computed this fraction by dividing cumulative sum of the principal component variances (i.e. eigenvalues) by the sum of the variances.

We plotted the first two components out of the twenty components (Figure S3b), the first one showed a smooth bump kernel and the second one showed a smooth primacy kernel. Together they count for 70% of the total variance explained, which indicates that the main factors contributing to kernel shape are not the high frequency oscillations but these low frequency kernel shapes.

**Figure S3:**
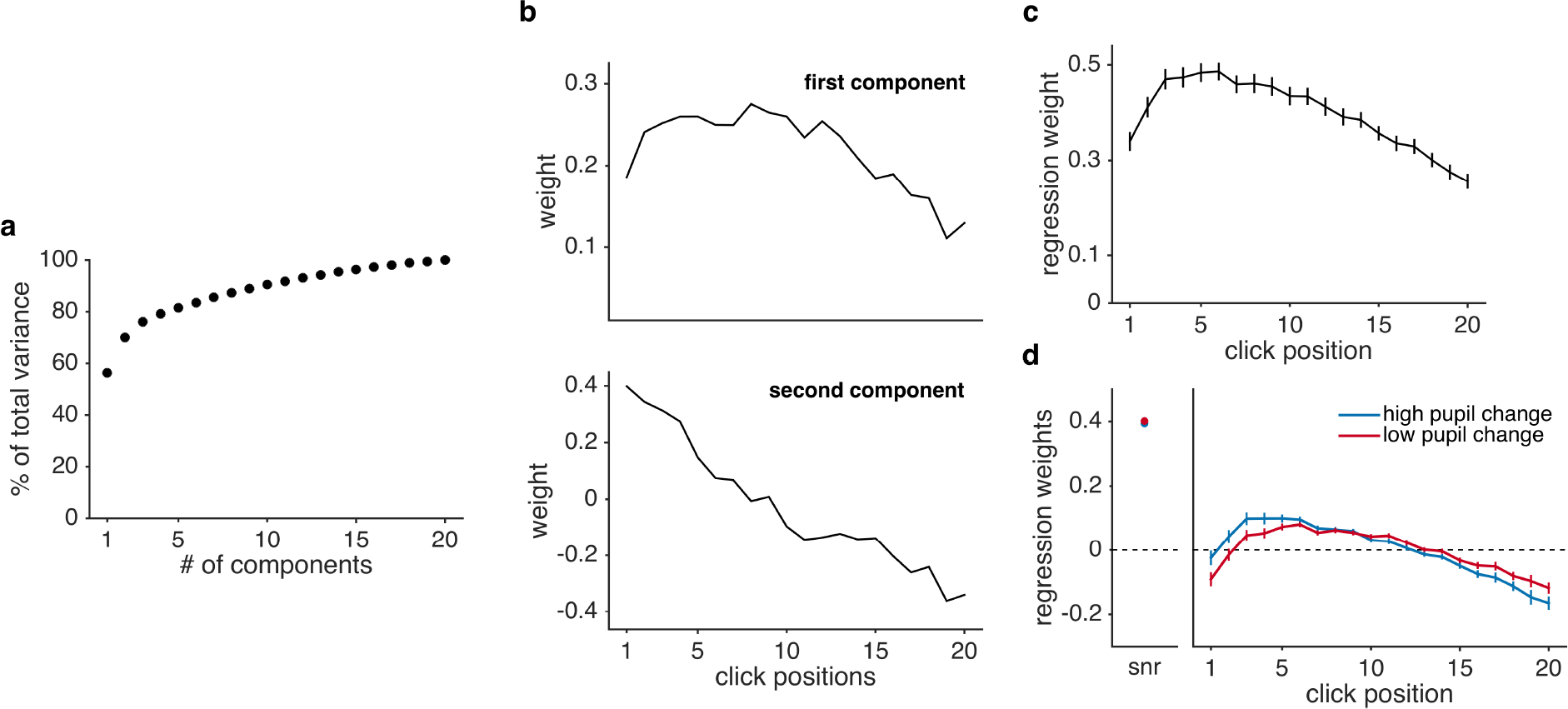
(a) Cumulative sum of fraction of total variance retained as the number of components increased. (b) First two principal components. (c) Integration kernel created using only the first two principal components. (d) Regression weights (recreated using only the first two principal components) averaged across participants split into high vs. low pupil change groups for visualization.

We performeddimensionalityreductiononthekernelweightsbypickingthesefirst two components to recreate the kernel weights (Figure S3c), and tested for the effect of interaction between pupil change and time on kernel weights. With two way mixed ANOVA, we saw a significant effect of interaction between pupil change and time on kernel weights (F(19,2014) = 4.096, p = 6.9e-9), which concur with what was reported in the original manuscript. This analysis was done with pupil change a continuous variable, but again for visualization purpose, we plot the kernel weights of participants split by high vs low pupil change (Figure S3d). These results combined suggest that the pupil effect on kernel shape is not due to the high frequency components of the kernel, but the low frequency components (the bump and the primacy kernels).

### S.4 Relationship between kernel shape and pupil measured using different window sizes (corresponding to Main Text Section 2.4)

One potential concern is that the window for measuring pupil change to the stimulus window (1 second after stimulus onset) is also a relatively early window for inspecting pupil change, and that this relatively early window can bias the result to detecting relationships between pupil and processes that happen earlier in time. We addressed this concern in two ways:

1. An expanding window analysis, in which we time-locked the pupil response to stimulus onset and expanded the window of analysis from between 0s and 1s to between 0s and 2s, and computed the pupil change as the difference between the maximum and the minimum of pupil response for each window size.
2. A sliding window analysis, in which we computed the pupil change as the difference between every point on the pupil response time course and baseline (.25 sec pre stimulus).

#### S.4.1 Expanding window analysis

We repeated the analysis with a two-way mixed ANOVA to quantify the effect of pupil on signal-to-noise ratio and on kernel shape. As in the original reported results, we did not see a significant influence of pupil on signal-to-noise ratio, but we did see a significant effect of pupil on integration shape at every window size (FDR corrected p-values for all points <= 0.011). We reported the corrected p-values of the interaction effect between pupil and time on kernel weights in Figure S4a.

To inspect whether the direction of this result was consistent with the reported result, and to inspect whether it was the same set of clicks (i.e. the clicks occuring early on in the stimulus) that contribute to the change in kernel shape, we looked at the correlation coefficients between pupil change (for every window size) and kernel weights. We found that, as with the original reported results, only the 2nd and 3rd kernel weights correlate significantly positively with pupil change across individuals (Figure S4b). This suggested that later pupil change did not reveal relationship with integration that happened later in time.

**Figure S4:**
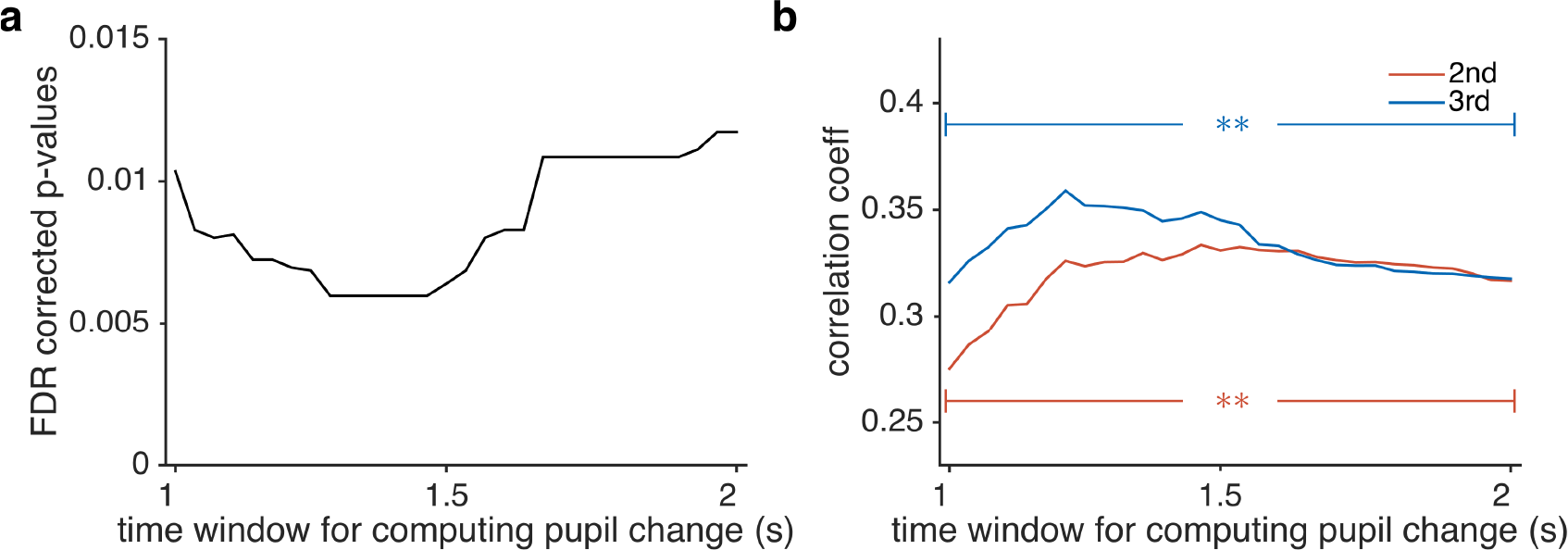
(a) FDR corrected p-values of effect of interaction between pupil and time on kernel weights. (b) All correlation coefficients are significantly positive (FDR corrected p-values for all coefficients <= 0.005)

#### S.4.2 Sliding window analysis

**Figure S5:**
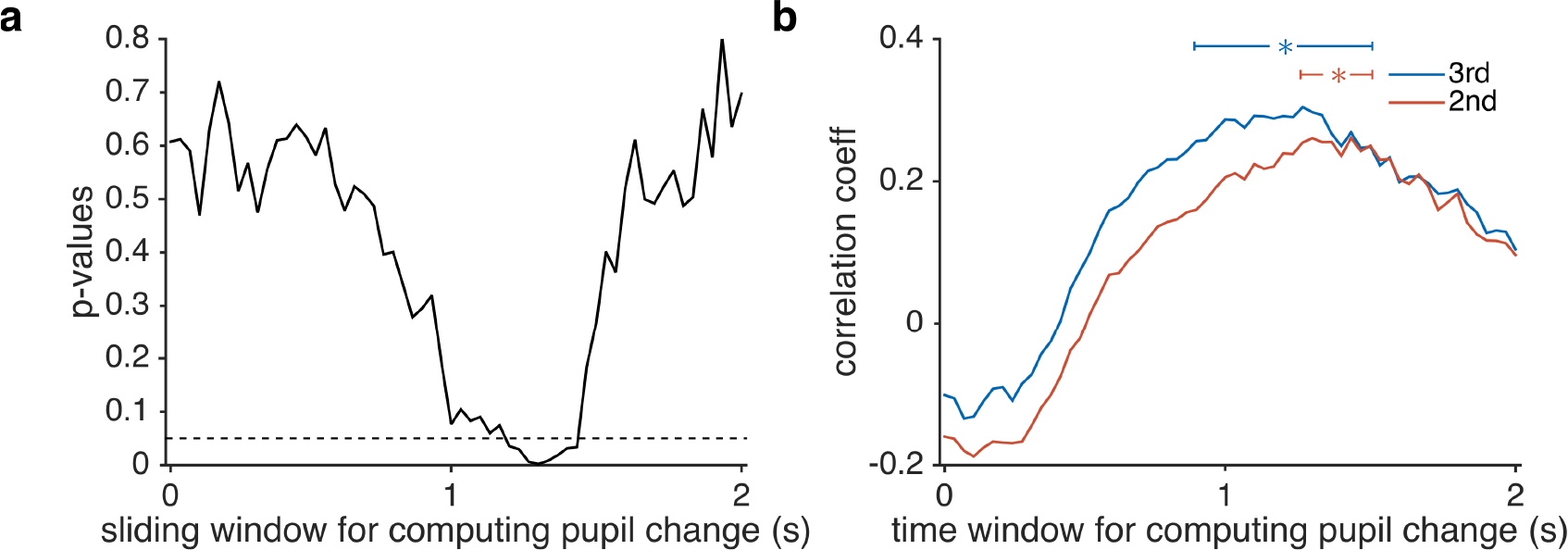
(a) Uncorrected p-values of effect of interaction between pupil and time on kernel weights. (b) Correlation coefficients between pupil and second/third kernel weights (*p* <= 0.01 uncorrected)

We repeated the above analysis with a sliding window analysis, and saw the similar results. Specifically, we used each point in time of the pupil response between 0 and 2 seconds after the clicks onset. Again a two way mixed ANOVA revealed that pupil change modulated kernel shape, although the largest effect was observed about 250ms after the end of the clicks (Figure S5a). In addition, we again saw that only the 2nd and the 3rd kernel weights correlated with pupil change (Figure S5b). Combined, these analyses support that the relationship between kernel shape and pupil is not a result of early window of pupil change biasing the detection of modulation in early kernel weights.

### S.5 Relationship between sequential effects and pupil measured using different window sizes (corresponding to Main Text Section 2.5)

In contrast to our results, Urai et al 2017 found an association between pupil response and the magnitude of the history effects. The question here, then, is whether such a pupil-history effect relationship exists in our data. One key difference between our analysis and Urai’s is that we only looked at pupil on the present trial, whereas Urai looked at pupil on the previous trial. To address this question comprehensively, we repeated the above sliding window analysis to inspect correlation between pupil on the previous trial and history effects at the within participant level.

#### S.5.1 Expanding window analysis for pupil on previous trial

Here we used a similar expanding window analysis as described in the previous section, and repeated the regression analysis described in Main Text Section 2.5 (equation (4)). The main difference was that the pupil signal comes from the previous trial, not the current trial. Repeating the regression analysis with these two measures we did not see a significant interaction effect between previous trial effect and pupil change (Figure S6) suggesting that the relationship between past pupil and history effects is not present in our data.

**Figure S6:**
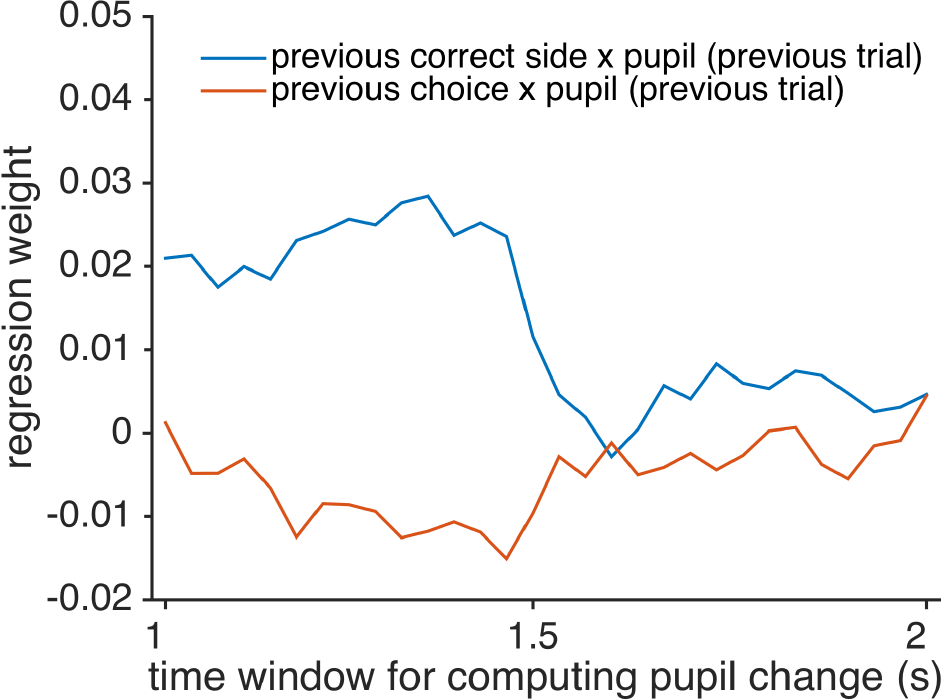
Expanding window analysis of interaction between previous trial effects and pupil change.

#### S.5.2 Sliding window analysis for pupil on previous trial

We also used a sliding window approach to investigate the interaction effect between previous trial and pupil change on the last trial. Again, we did not observe any significant effect (Figure S7), further confirming the lack of a relationship between pupil and history effects in our data.

Clearly, these null results are quite different from the findings of Urai et al. and the obvious question is why this difference arises. As discussed in the Discussion, we believe there are two possible issues at play here. First, the history effects we observed in this task are relatively small (having the same effect on choice as approximately 1 click). This small effect size for past trials means that the modulation of this small effect by pupil will be difficult to detect. The second reason we may not have observed the effect is down to the timing of our trials. To maximize the number of trials in the task, the inter-trial interval was short. This short ITI makes it hard to examine the late pupil components, which is exactly the component that Urai et al. found correlating with history effects.

**Figure S7:**
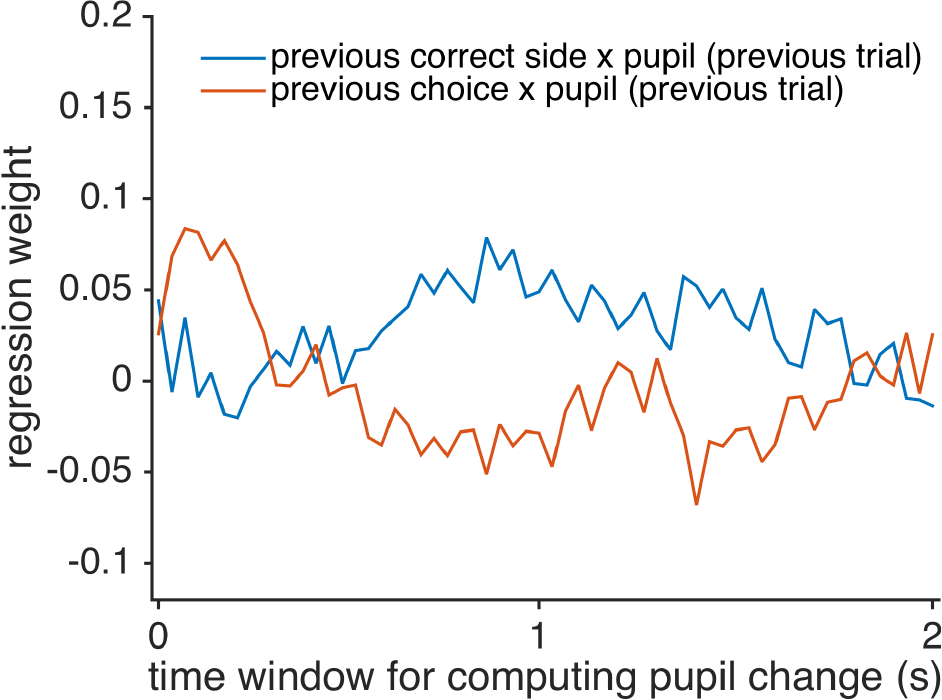
Sliding window analysis of interaction between previous trial effects and pupil change.

### S.6 Variance inflation factor analyses for regression models

Most of the reported analyses require fitting a considerable number of regressors, which raises the concern that the regressors may be collinear with each other. Here we computed variance inflation factors (VIFs) for our regression models to examine if the regressors in our regression models are collinear with each other.

For equation (3), we note that VIFs are all around 1 (the minimum value for variance inflation factors) for all participants (Figure S8), and thus it does not run into multicollinearity issues.

**Figure S8:**
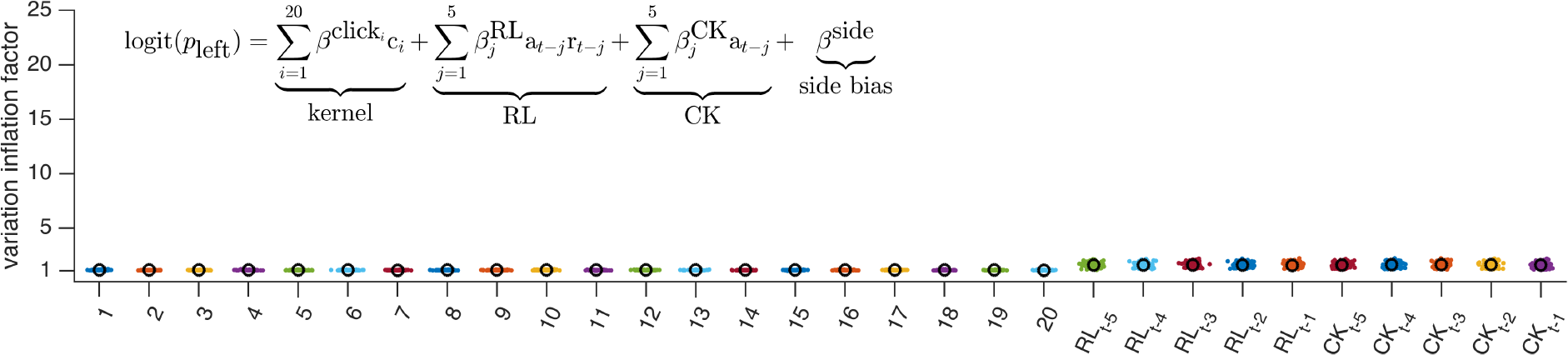
VIFs for equation (3). Equation replicated in figure. Each participant’s VIF is shown as a colored dot, the mean across participants is shown as a black circle.

Beginning with equation (4) (which had fewer regressors than equation (5), namely trial-by-trial analysis showing interaction effect between pupil and signal-to-noise ratio),we find that VIFs for individual clicks (1 to 20) and side × pupil regressors are low (Figure S9). Mean VIFs for all the other regressors are <= 10, which is the standard cutoff for diagnosing multicollinearity in regression [47, 48].

**Figure S9:**
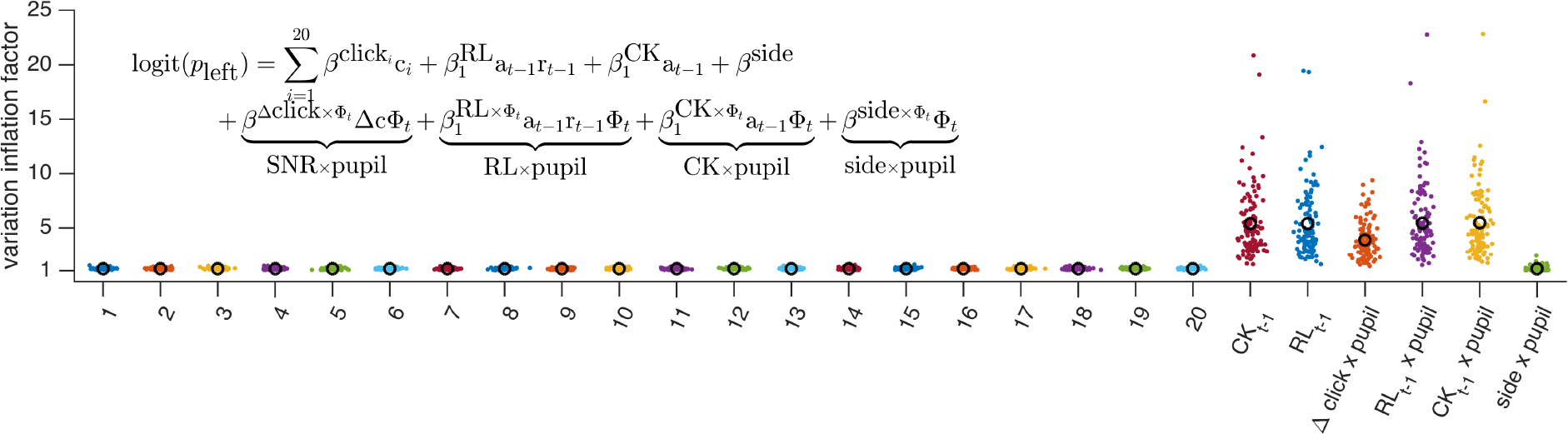
VIFs for equation (4).

However, we see some participants with VIFs much higher than 5. To assuage the validity of this regression analysis, we excluded these participants and picked only participants for whom all the regressors have a VIF smaller than or equal to 5, and ran the regression analyses on only these participants (n = 54). We plot the VIFs from the new group of participants (Figure 8). In the new group of participants, we see the same results as reported in Main Text Figure 5): trial-by-trial analysis of interaction effect between pupil and signal-to-noise ratio is significantly negative (Beta = −0.0083, FDR corrected for multiple comparisons p = 0.01) (Figure 9).

**Figure S10:**
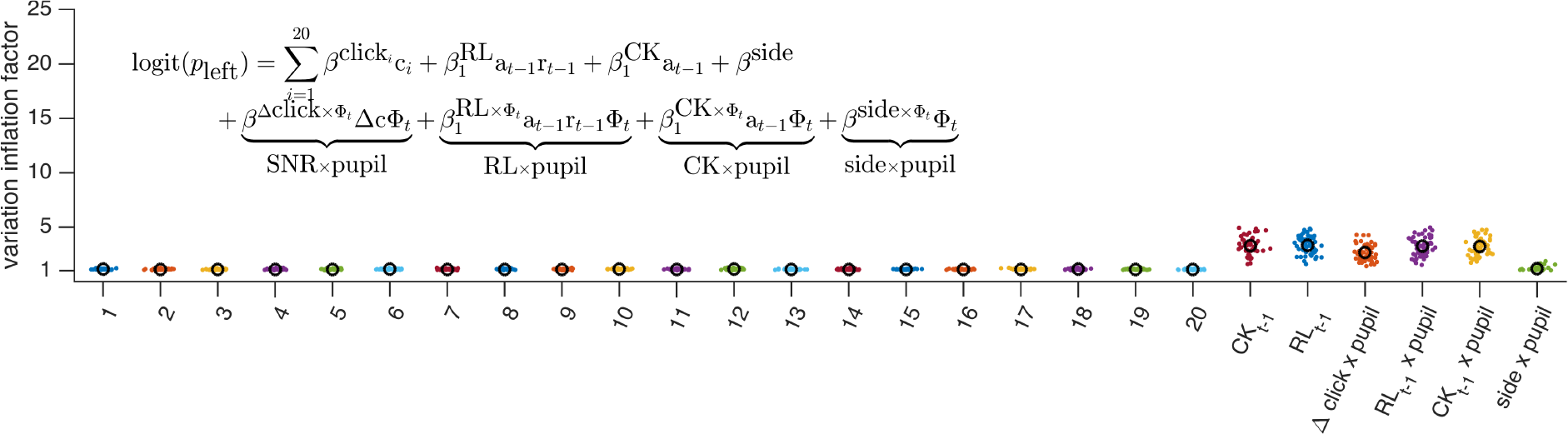
VIFs of only participants for whom all the VIFs are smaller than or equal to 5 (n = 54) for equation (4)

**Figure S11:**
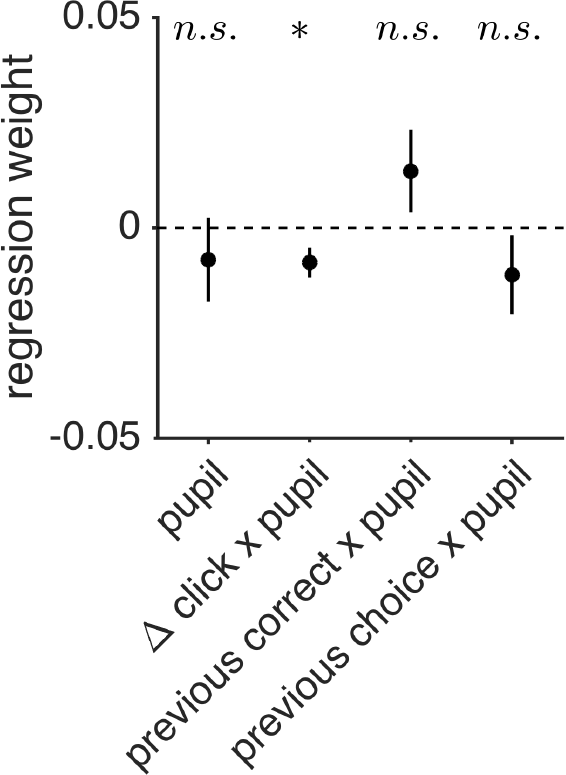
Trial by trial analysis of interaction effects between pupil and behavior with only participants for whom all the VIFs are smaller than or equal to 5 (n = 54).

For equation (5), due to the large number of regressors, the VIFs for all regressors are relatively high (Figure S12). We acknowledge that for this specific regression model (inspecting pupil interaction with kernel shape on a trial by trial basis) we are limited in what we can do with the amount of data we have. However, the mean VIFs for all regressors are still smaller than 10, which is not a definitive diagnosis that the model has serious multicollinearity issues [47].

**Figure S12:**
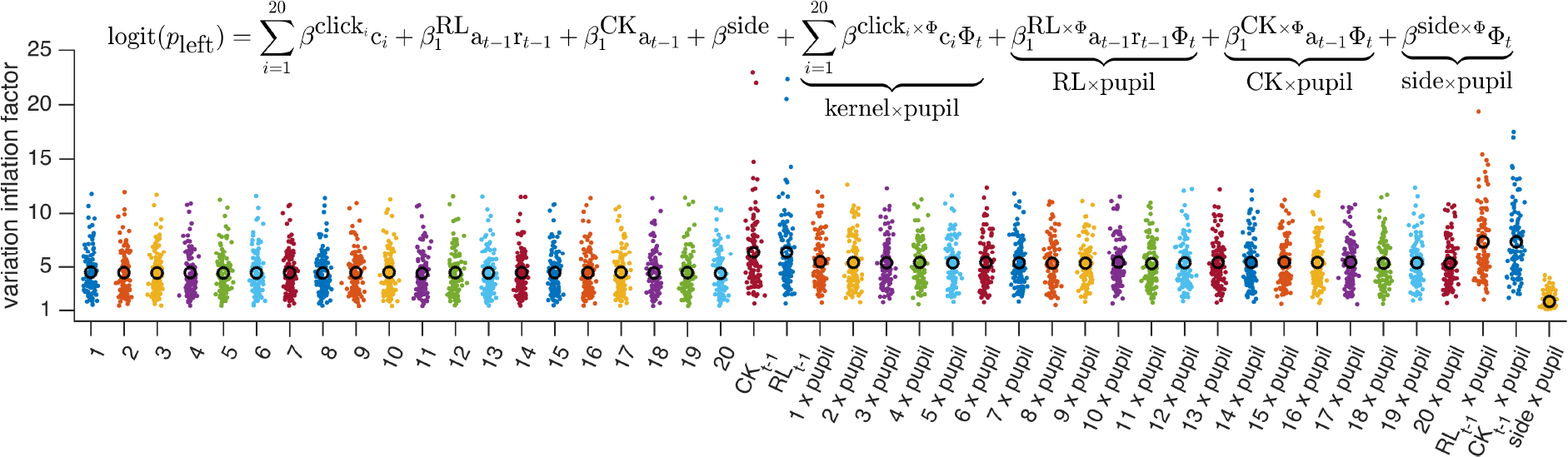
VIFs for equation (5)

### S.7 Pupil analysis using pupil change residualized from previous trial pupil change

The ITI is relatively short for pupil signal, and one potential concern is a bleed over effect of pupil signal from the previous trial into the current trial. To alleviate this concern, we repeated our analyses with residualized pupil change. We took the regression of the pupil change in the previous trial on pupil change in the current trial, and used the residual as our new measure of pupil change. Repeating the trial-by-trial analyses, we see that our results concur with the original results. Specifically we find that Δ*click* × *pupil* has a significant interaction effect on choice (Beta = −0.0184, FDR corrected p = 0.0025)(Figure S13).

**Figure S13:**
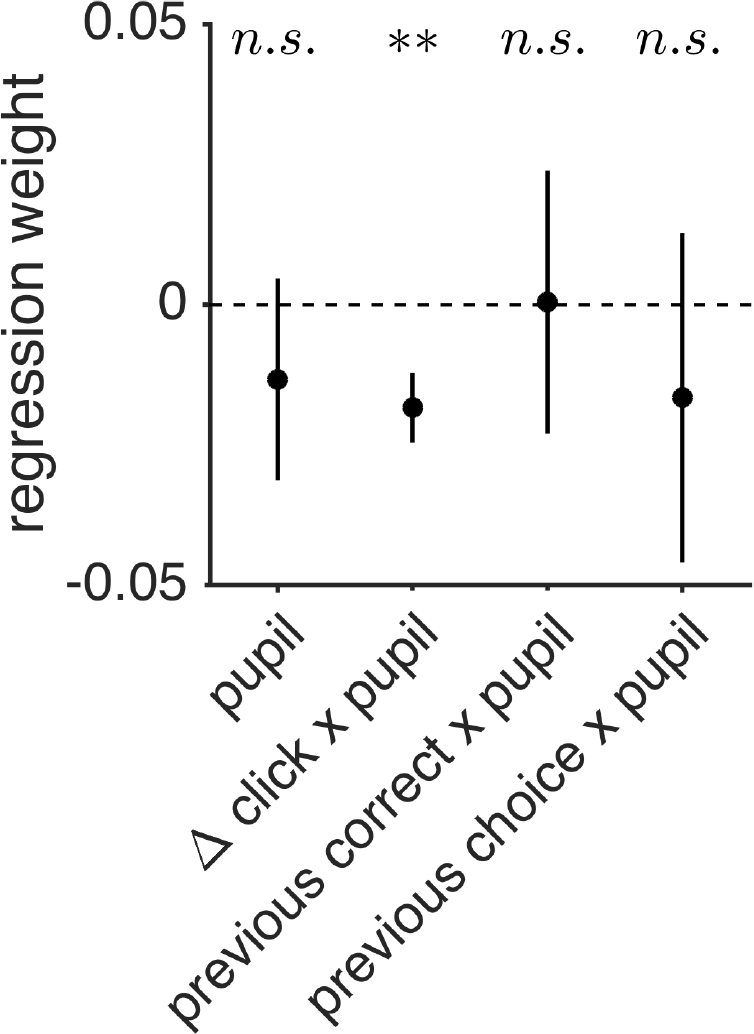

### S.8 Categorizing kernel shapes

To categorize the kernel for each participant into one of the four shapes, we fit polynomial functions with different degrees to participants’ choices, and selected the best fitting model with model comparison using the Akaike Information Criterion (AIC). Specifically, we assume that the probability of choosing left at trial *t* is (the logit of) the weighted sum of clicks, where the weights are from a polynomial function, as shown in the following equation:

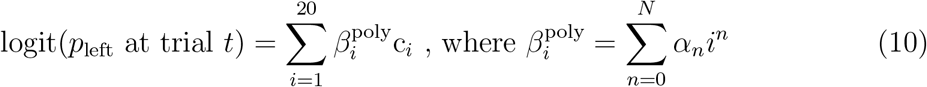

We fitted three different polynomial functions by changing N from 0 to 2: constant, linear, and quadratic. We then selected the best fitting function for each participant by comparing the fits from different polynomials with AIC. We categorized each participant’s integration kernel into one of the four shapes using the following criteria: (1) flat: kernel was best fit with the constant function; (2) primacy: kernel was best fit with linear function with a negative slope (*α*_1_), or with quadratic function with a minimum (*α*_2_ > 0) and the minimum is located later than the 10th click, or with quadratic function with a maximum (*α*_2_ < 0) and the maximum is located earlier than the 2nd click; (3) recency: kernel was best fit with linear function with a positive slope, or with quadratic function with a minimum (*α*_2_ > 0) and the minimum is located earlier than the 10th click, or with quadratic function with a maximum (*α*_2_ < 0) and the maximum is located later than the 18th click; (4) bump: kernel that did not meet the previous three criteria (i.e. kernel was best fit with quadratic function and was neither primacy nor recency).

**Figure S14:**
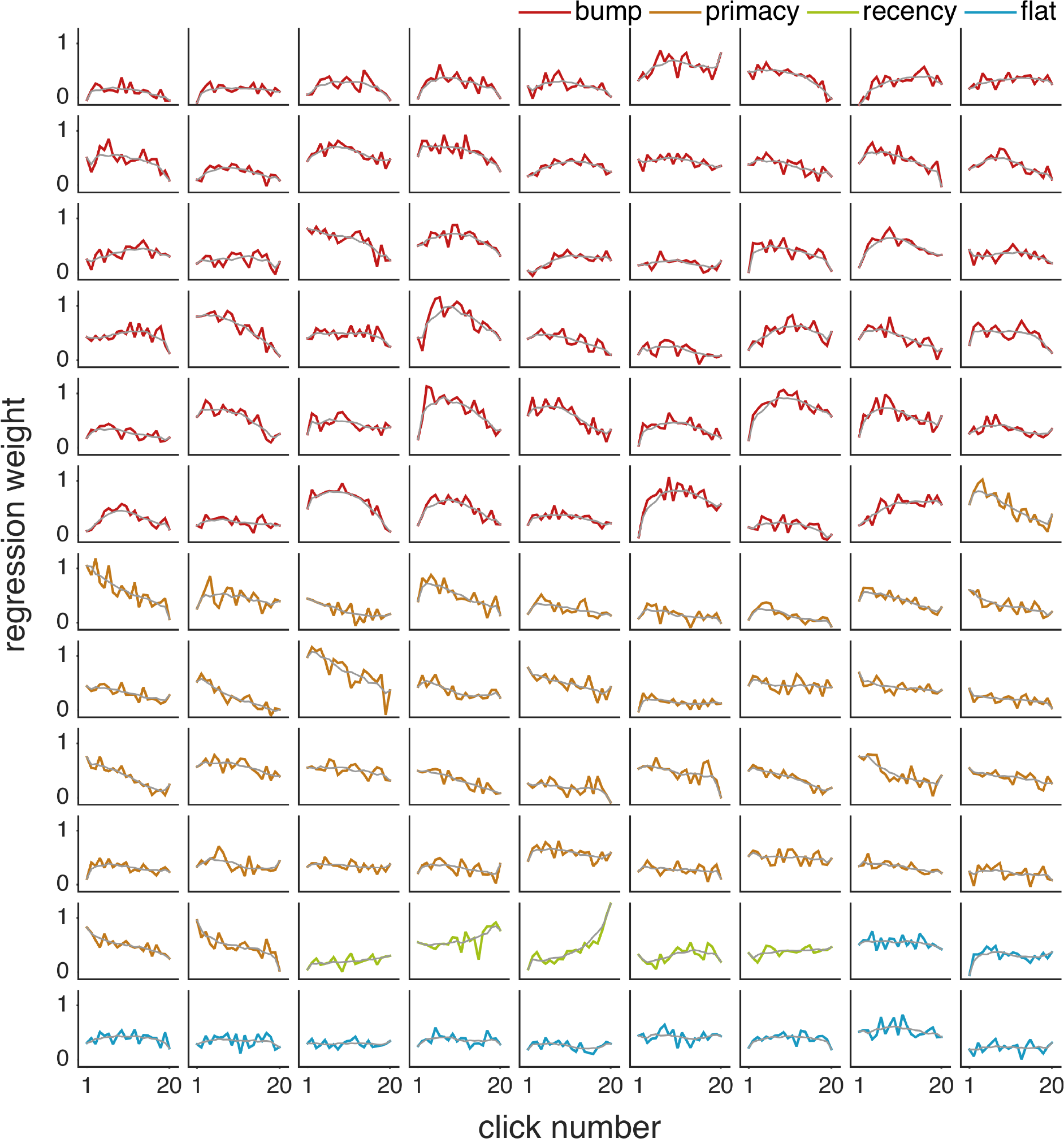
Individual integration kernel plots. Colored lines are regression weights of clicks from equation (3). Plots are sorted and color coded by kernel shape. Light grey line shows smoothed integration kernel.

